# Comparison of CPU and GPU Bayesian Estimates of Fibre Orientations from Diffusion MRI

**DOI:** 10.1101/703835

**Authors:** Danny H.C. Kim, Lynne J. Williams, Moises Hernandez-Fernandez, Bruce H. Bjornson

## Abstract

**Background:** The correct estimation of fibre orientations is a crucial step for reconstructing human brain tracts. Bayesian Estimation of Diffusion Parameters Obtained using Sampling Techniques (*bedpostx*) is able to estimate several fibre orientations and their diffusion parameters per voxel using Markov Chain Monte Carlo (MCMC) in a whole brain diffusion MRI data, and it is capable of running on GPUs, achieving speed-up of over 100 times compared to CPUs. However, few studies have looked at whether the results from the CPU and GPU algorithms differ. In this study, we compared CPU and GPU *bedpostx* outputs by running multiple trials of both algorithms on the same whole brain diffusion data and compared each distribution of output using Kolmogorov-Smirnov tests.

**Results:** We show that distributions of fibre fraction parameters and principal diffusion direction angles from *bedpostx* and *bedpostx_gpu* display few statistically significant differences in shape and are localized sparsely throughout the whole brain. Average output differences are small in magnitude compared to underlying uncertainty.

**Conclusions:** Despite small amount of differences in output between CPU and GPU *bedpostx* algorithms, results are comparable given the difference in operation order and library usage between CPU and GPU *bedpostx*.

## Background

Recent human brain MRI data sizes are continuing to increase as large imaging data repositories curate structural, functional and diffusion data with higher spatial resolution, faster temporal sampling, and higher angular (Van Essen et al. 2013; Sotiropoulos et al. 2013; Miller et al. 2016). Performing image analysis on such dataset is computationally expensive, even with clusters of CPUs working simultaneously on a single dataset (Hernandez-Fernandez et al. 2019). By contrast, graphics processing units (GPUs) have a massively parallel structure designed with hundreds of smaller cores optimized to exploit the data level parallelism of certain applications, utilizing simpler instruction sets and distributing them over multiple cores (Eklund et al. 2013a; Hernandez-Fernandez et al. 2013). This parallelization can accelerate computationally slow processes such as data visualization, stochastic iteration, and Bayesian simulations including probabilistic tractography (Chang et al. 2014; Eklund et al. 2013a; Eklund et al. 2013b; Hernandez-Fernandez et al. 2013; Hernandez-Fernandez et al. 2019; Lee and Kim 2013; McGraw and Nadar 2007; Sotiropoulos et al. 2013). A popular tool in estimating diffusion parameters for whole brain diffusion MRI is available to be run on both CPU or GPU, with GPU algorithm achieving over 100 times speed-up compared to its CPU algorithm (Behrens et al. 2007; Behrens et al. 2003; Hernandez-Fernandez et al. 2013). Despite the GPU’s advantages in acceleration, few studies have examined whether there are differences in computational output from the CPU and GPU. In general, checking for output convergence between CPU and GPU results is important for several reasons. First, CPU and GPU both have double-precision capabilities in their compilation and runtime libraries, but the optimization of performance and speed-up of GPU binaries may restrict them to using single-precision libraries which can cause results to be different due to float-point precision differences (Colberg and Hofling 2011; Whitehead and Fit-Florea 2011). Secondly, there are differences in the CPU and GPU random number generators and operation orders in implementing Markov Chain Monte Carlo (MCMC) (Hellekalek 1998; Luizi et al. 2010; Park and Miller 1988). For GPU results to be used interchangeably with existing CPU algorithms, the GPU algorithm should produce results that are reproducible and convergent with results obtained by the CPU algorithm. For example, Hernandez-Fernandez et al., compared the mean of a few representative diffusion weighted voxel values in a repeated test between CPU and GPU and found almost identical results (Hernandez-Fernandez et al. 2013). However, their study did not report on CPU/GPU differences in contiguous within-slice voxels or multi-slice brain data. The current study aims to extend these findings by comparing sampled distribution shapes of CPU and GPU Bayesian estimation of diffusion parameters in a whole brain dataset (Behrens et al. 2007; Behrens et al. 2003; Hernandez-Fernandez et al. 2013).

This paper is organized as follows. Brief introductions of DTI and Bayesian estimation of diffusion parameters are given. Then, the complete methodology of output comparison technique is described. Results of output comparison are presented for each diffusion parameter type, and then, we give our conclusions and discussions

### Diffusion MRI and Bayesian Estimation

Diffusion MRI (dMRI) is a useful tool in visualizing the white matter connectivity of the brain and is widely used in both research and clinical contexts. dMRI is sensitive to molecular diffusion of water and enhances the anisotropy—the directional dependence—of neuronal white matter fibre tracts, which can be used to create fractional anisotropy maps, mean diffusivity maps and fibre pathways (Beaulieu 2002; Johansen-Berg and Rushworth 2009). A commonly used method to estimate the fibre orientations and reconstruct the brain tracts in vivo is to use the FMRIB Software Library’s (FSL) “Bayesian estimation of diffusion parameters” (*bedpostx*) and “probabilistic tracking of crossing fibres” (*probtrackx*) algorithms. In brief, *bedpostx* employs a Markov Chain Monte Carlo sampling technique to estimate the posterior probability density functions (PDF) of the diffusion parameters utilizing the “ball-and-stick” model which takes into account multiple fibre orientations in a given voxel where appropriate (Behrens et al. 2007; Behrens et al. 2003). This allows the resolving of within-voxel fibre crossings, which is a common hurdle during the fibre tracking step, by fitting more than one fibre orientation in a given voxel only when it is relevant to do so. This feature is the “automatic relevance determination” (ARD) algorithm in *bedpostx* (Behrens et al. 2007) which initially sets the additional fibre fractions in secondary orientations to zero with low variance, and iteratively estimates the variance separately so that when the additional fibre orientation is supported by the data, the additional fibre fraction can take a non-zero value with a larger variance. *bedpostx* uses the Levenberg-Marquardt (L-M) fit to initialize parameters by minimizing the sum of squared model residuals, similar to fitting a diffusion tensor model, then, it proposes a value for each parameter, drawing from Normal proposal distribution, calculates the likelihood term, and accepts or rejects the proposed value based on a Metropolis acceptance criterion. When employing the ball-and-stick model where the isotropic compartment is fitted with a mean value within a voxel (i.e. model=1), *bedpostx* gives the following PDF distributions for each voxel as output: diffusivity value (*d*), baseline signal (*S0*), weight of each fibre orientation’s contribution to anisotropic diffusion signal (stick), also known as fibre fraction values (*f*_1_, *f*_2_, etc.), and each fibre orientation’s directional angles expressed in polar coordinates (ϕ_1_, θ_1_, ϕ_2_, θ_2_, etc). These PDFs are then randomly sampled by *probtrackx* to create fibre streamlines through stochastic propagation of multiple particles through the diffusion space (Behrens et al. 2007; Behrens et al. 2003). Because *bedpostx* processes each voxel serially in the CPU, an extensive amount of computational time is required to obtain the PDFS, which makes it impractical for utilization in a clinical medical environment where a computing cluster may not be available (Lerner et al. 2013; Yamada et al. 2009). To alleviate this problem, and to reduce computational time substantially FSL provides a GPU-based parallelized version of *bedpostx*, called *bedpostx_gpu* (Hernandez-Fernandez et al. 2013). Here, the L-M initialization and MCMC sampling are parallelized such that multiple voxels are processed simultaneously. Difference in operation order exists between *bedpostx* and *bedpostx_gpu* such that, in the GPU, L-M initialization for the entire brain is done first, then MCMC sampling are done for the entire brain, whereas in the CPU, L-M initialization and MCMC sampling are done in sequential order for each voxel. We know of no study to date that has quantitatively examined output similarities and differences between the *bedpostx* and *bedpostx_gpu* algorithms in a whole-brain DTI dataset. Further, because the PDF distributions obtained from *bedpostx* is used in obtaining fibre streamlines in *probtrackx* and not their mean values, differing distributional shapes between the two algorithms can also cause bias in output fibre tracking using *probtrackx*. This study aims to compare the output of *bedpostx* and *bedpostx_gpu* and report on output PDF distribution (*f*_1_, *f*_2_, ϕ_1_, θ_1_, ϕ_2_, θ_2_) shape difference, magnitude of difference in mean value and underlying uncertainty value.

## Methods

### A. Computational Resources

*bedpostx* was used for output comparison with the GPU version *bedpostx_gpu*, both from FSL 5.0.8 package running on Ubuntu 10.04 LTS. The CPU version ran on a workstation with a dual Intel Xeon X5670 2.93 GHz CPU with 6 x 4-GB DDR3-1333 memory, and 24 threads. The GPU version ran on a workstation with one NVIDIA Tesla C2075 with 448 CUDA cores, 6-GB GDDR5 dedicated memory, PCIe x16 bus, CUDA 5.5 with driver version 331.75.

### B. Data

Diffusion data from the Human Connectome Project (HCP) database (subject: mgh1005) was used for running multiple trials of *bedpostx* and *bedpostx_gpu*. The full dMRI data consist of directional volumes acquired in multiple shells (b=0,1000,3000,5000,10000) but for our work, a single shell from the full set was used for analysis: motion and eddy corrected, b=1000, 64 directional volumes and 6 non-directional volumes, 1.5mm isotropic, 140×140×96. This was chosen because most clinical and research studies have access to a similar single-shell dMRI acquisition method and the resulting data can still support multi-fibre modeling of *bedpostx* algorithms. T1-weighted anatomical scan of the same subject was segmented (Zhang et al. 2001) to derive binary masks of grey matter, white matter and cerebrospinal fluid, then co-registered to the diffusion data (Jenkinson et al. 2002; Jenkinson and Smith 2001). These masks were used to quantify how many significantly different distributions were localized in each tissue class.

### C. Bedpost PDF creation

Specified *bedpostx* and *bedpostx_gpu* input parameters are: 2250 MCMC iterations, of which during the latter 1250 iterations, parameter values were recorded to PDF every 25 iterations, resulting in 50 samples per PDF; monoexponential model (i.e. fit with mean diffusivity) with ARD fitting 2 fibres per voxel where appropriate. 20 trials of *bedpostx* and *bedpostx_gpu* were run with different random number generator seeds and output distributions from all trials were merged together to form 1000 samples per parameter PDF for *bedpostx* and for *bedpostx_gpu*. Furthermore, to inspect differences in L-M initialization between *bedpostx* and *bedpostx_gpu*, 20 trials of each algorithm were run again but with 1 iteration to record 1 sample close to the initializing value.

### D. PDF distribution comparison and statistical analysis

PDF shape was statistically compared via two-sample Kolmogorov-Smirnov (KS) test to derive voxels that have different distributions between CPU and GPU (two-tailed, p < 0.05, uncorrected). Family-wise error rate was controlled by the Bonferroni method (Holm 1979). Voxels with significantly different distributions were then further categorized by their KS-scores (S) in 4 different ranges: 0.1-0.2, 0.2-0.3, 0.3-0.4, > 0.4. S scores illustrate the amount of sample deviation (e.g. S = 0.35, 35% of samples differ between two distributions). For *f*_1_ and *f*_2_, CPU mean values along with absolute difference in mean CPU/GPU values were calculated and averaged for each S range. For angles, mean, standard deviation and median difference in principal diffusion directions (PDD) along with 95^th^-percentile cones of angular uncertainties (CAUs) were calculated in voxels with at least one significantly different angle parameter for each pairs (i.e. ϕ_1_ OR θ_1_, ϕ_2_ OR θ_2_). Maximum S score between the [ϕ,θ] pair was used when categorizing significantly different angle parameters into different S-ranges.

### E. Effect of mixed fibre fraction and orientation samples near crossing-fibre areas

Fibre fraction parameters (*f*_1_, *f*_2_, etc.) and their associated orientations (ϕ_1_θ_1_, ϕ_2_θ_2_, etc.) could be inconsistently associated with the different underlying sub-fibre populations, especially if the fibre fractions are of comparable strength (Jbabdi et al. 2010). This can cause differing proportions of fibre fraction and orientation values to be labeled as one group (e.g. *f*_1_,ϕ_1_θ_1_) but labeled as another on the next trial (e.g. *f*_2_,ϕ_2_θ_1_). There is no guarantee that the labeling happens consistently and because we are merging samples from 20 different *bedpostx* and *bedpostx_gpu* trials to form the PDF distributions for comparison, it is possible that differences between the two platforms occur due to the this inconsistent labeling of sub-fibre populations. To investigate this effect of mixed fibre fractions and how much it may contribute to CPU and GPU output differences, we swapped *f*_1_,ϕ_1_θ_1_ and *f*_2_,ϕ_2_θ_2_ where *f*_2_ > *f*_1_ and ran the same statistical analysis on the swapped samples and compared the results against statistically different unswapped samples.

## Results

### A. Difference in L-M initialization

L-M initialization difference map is shown in Fig. 1 with difference greater than 0.5% or 1° of mean CPU values color coded. Diffusivity and baseline signal (*d, S0*) have increased difference towards the center of the brain, and *f*_2_, ϕ_2_, and θ_2_ have greater amount of different voxels compared to *f*_1_, ϕ_1_ and θ_1_.

**Figure 1:**
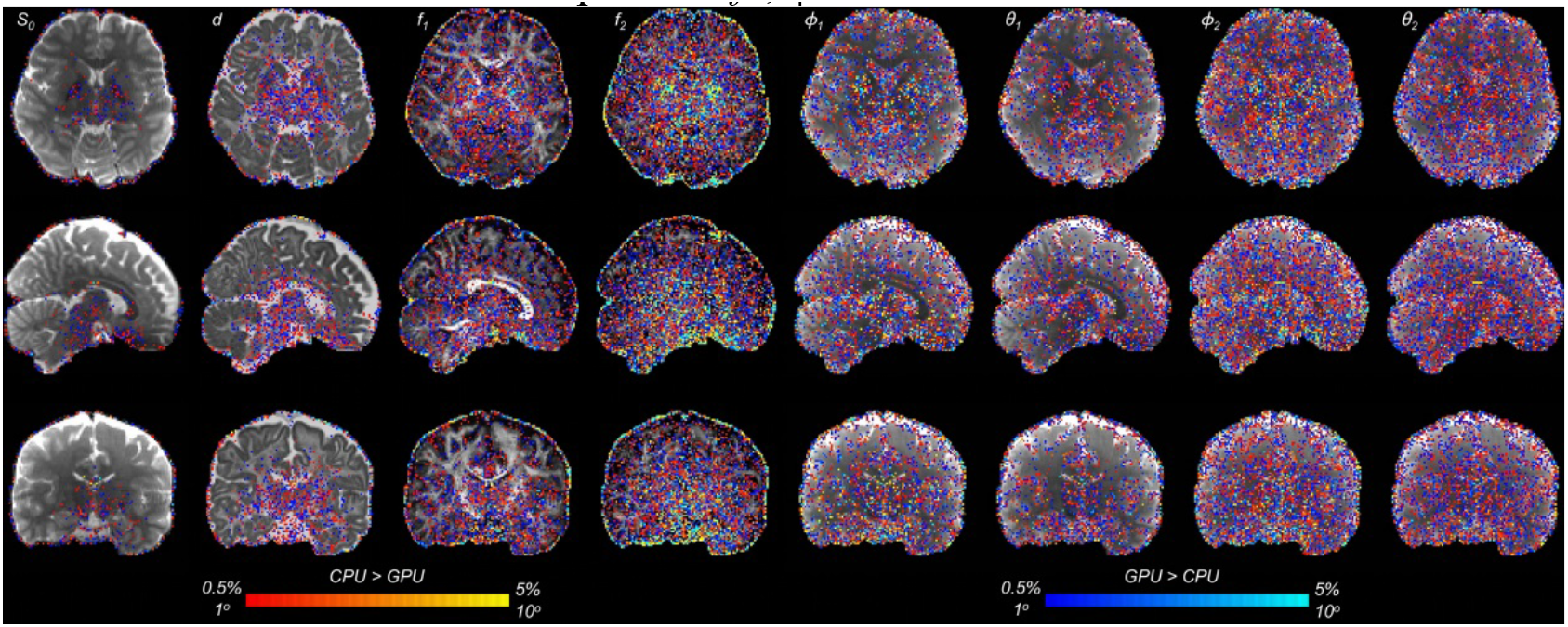
L-M initialization difference between CPU and GPU. Orange-yellow colors are CPU > GPU regions, and blue-light blue colors are GPU > CPU regions. Difference in scalar maps are thresholded at a magnitude of 0.5% with respect to mean CPU values and directional maps are thresholded at 1 degrees.

### B. *f*_1_

About 1% of total number of brain voxels (5139 of 436738) had significantly different *f*_1_ distributions. Significantly different voxels were sparsely localized throughout the brain bilaterally. Of the significantly difference voxels, 4% were found in cerebrospinal fluid, 29% were found in grey matter and 67% were found in white matter. The latter were located in long white matter projections, such as corpus callosum, corona radiata, internal capsules and anterior and posterior thalamic radiations (Fig. 2). Number of significant voxels, mean CPU *f*_1_, absolute average differences in mean *f*_1_, and absolute average difference in L-M initialization in each S-score region are summarized in Table 1.

**Figure 2:**
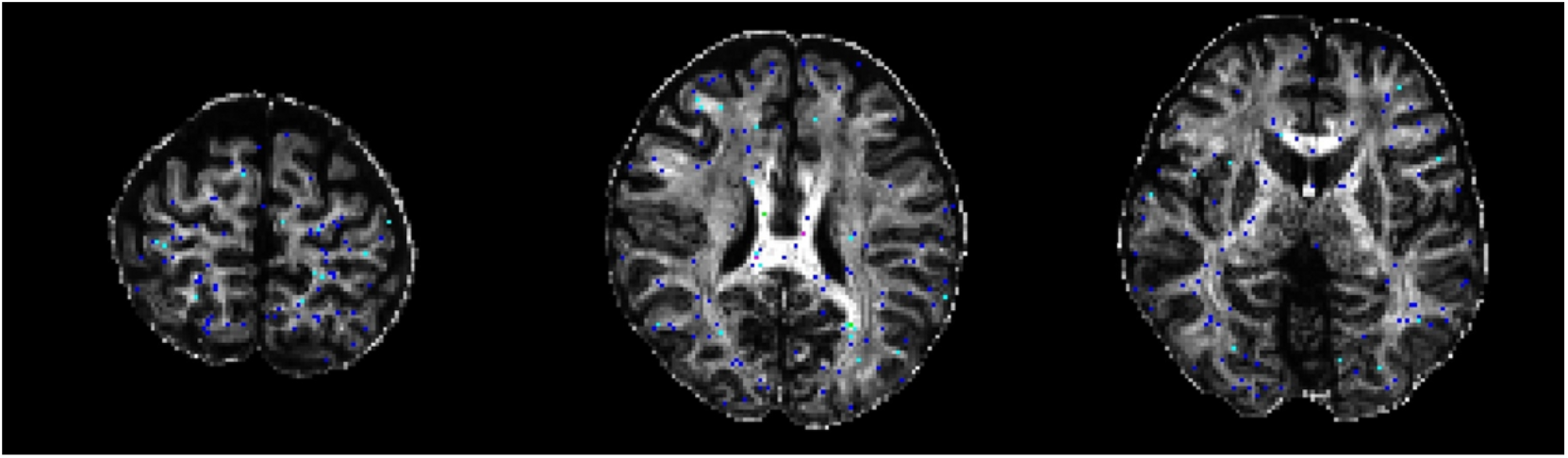
Significantly different *f*_1_ overlaid on mean *f*_1_ image. S-score ranges are in: 0.1-0.2=blue, 0.2-0.3=light blue, 0.3-0.4=green, > 0.4=magenta

**TABLE 1:** Significantly different *f*_1_ distributions: for each S-score range, averaged mean *f*_1_ of CPU distributions, averaged absolute difference in mean *f*_1_ and averaged absolute difference in L-M initialization are tabulated

| *S score*<br>(# of<br>voxels) | *Average*<br>*Mean $f_1$*<br>*CPU*<br>(*stdev*) | *Average*<br>*\|Mean $f_1$ CPU - Mean $f_1$*<br>*GPU\|* | *Average \|L-M CPU - L-M GPU\|* |
| --- | --- | --- | --- |
| 0.1 – 0.2<br>(4488) | 0.3262<br>(0.1613) | 0.0177 | 0.0008 |
| 0.2 – 0.3<br>(614) | 0.3964<br>(0.1505) | 0.0332 | 0.0009 |
| 0.3 – 0.4<br>(35) | 0.4707<br>(0.1786) | 0.0638 | 0.0206 |
| > 0.4<br>(2) | 0.7399<br>(0.1245) | 0.1333 | 0.2416 |
| *All* | 0.3357<br>(0.1623) | 0.0199 | 0.0010 |

Majority of voxels had S-scores less than 0.3 (5102 out of 5139). Example PDF distribution shapes of CPU and GPU in significant voxels derived from a single slice are shown in Fig 3. The largest S-score of 0.503 was found in the body of corpus callosum. Here, both *f*_1_ distributions have peaks near 0.99. GPU data had a sharper peak, with almost all samples above 0.9; whereas only half of CPU samples are above 0.9, with the remainder between 0.3 and 0.8. Here, average *f*_1_ initialization across 20 trials by CPU L-M was 0.52 while average *f*_1_ initialization by the GPU L-M across 20 trials was 0.99. Larger average L-M initialization differences were noted for larger S-score. After adjusting for *f_2_>f_1_* samples by swapping, 1513 voxels were no longer significantly different.

**Figure 3:**
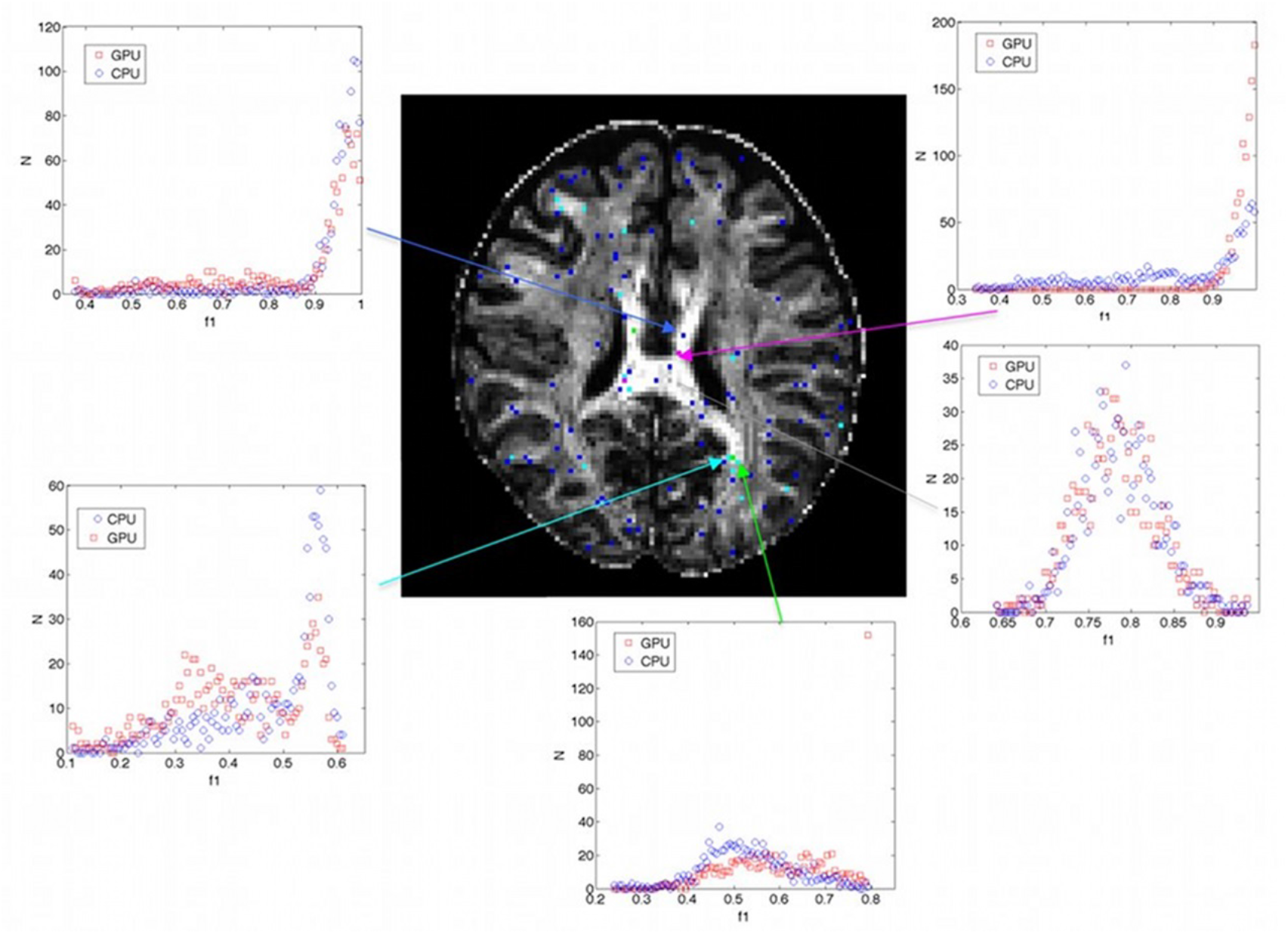
Example PDF distribution shape differences of significantly different *f*_1_. Grey arrow denotes non-significant distribution. Different ranges of S scores are depicted by following colors of arrows: blue 0.1-0.2, light blue 0.2-0.3, green 0.3-0.4, magenta >0.4

### C. *f*_2_

31% of total number of brain voxels (137061 out 436738) had significantly different *f*_2_ distributions. Significantly different distributions were localized in grey matter (44%), cerebrospinal fluid (33%) and white matter (23%). For the white matter, they were localized in long white matter projections similar to those identified in *f*_1_ (Fig. 4).

**Figure 4:**
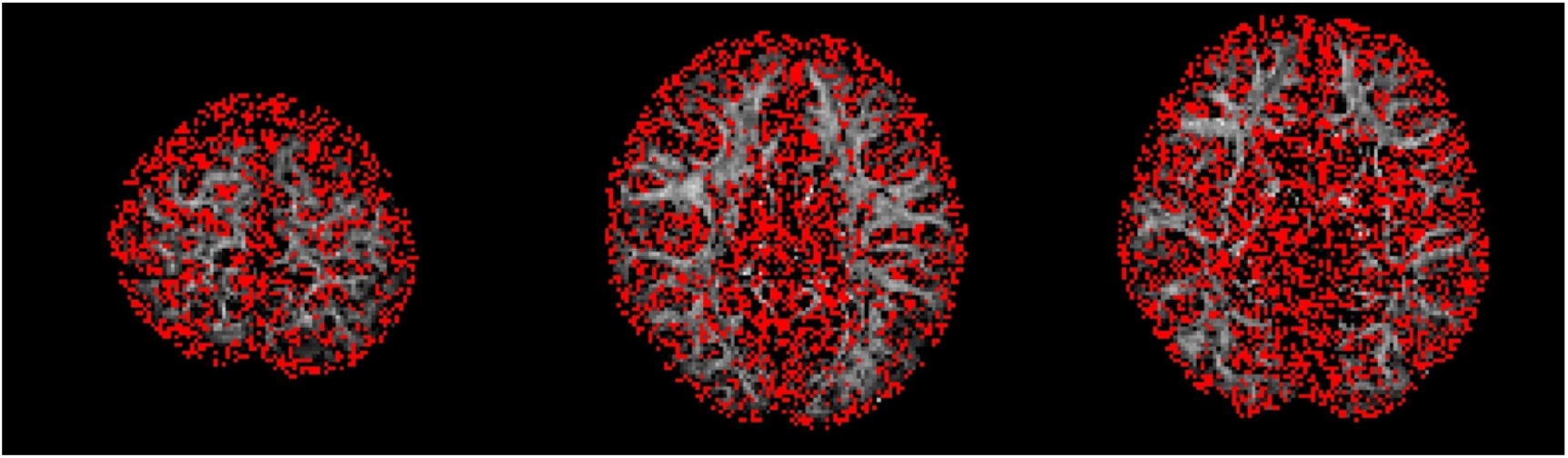
Significantly different *f*_2_ (red) overlaid on mean *f*_2_ image. Majority of voxels are located in grey matter and cerebrospinal fluid

85% of significant voxels had CPU or GPU mean *f*_2_ values lower than 0.05, predominantly in areas with grey matter and cerebrospinal fluid, likely the effect of ARD estimating *f*_2_ to zero in both *bedpostx* and *bedpostx_gpu*. To focus analysis on areas where *f*_2_ is supported by data, we reported mean CPU *f*_2_, absolute average differences in mean *f*_2_, and absolute average difference in L-M initialization in each S-score regions only on areas with mean *f*2 from CPU or GPU greater than or equal to 0.05 (Table 2).

**TABLE 2:**
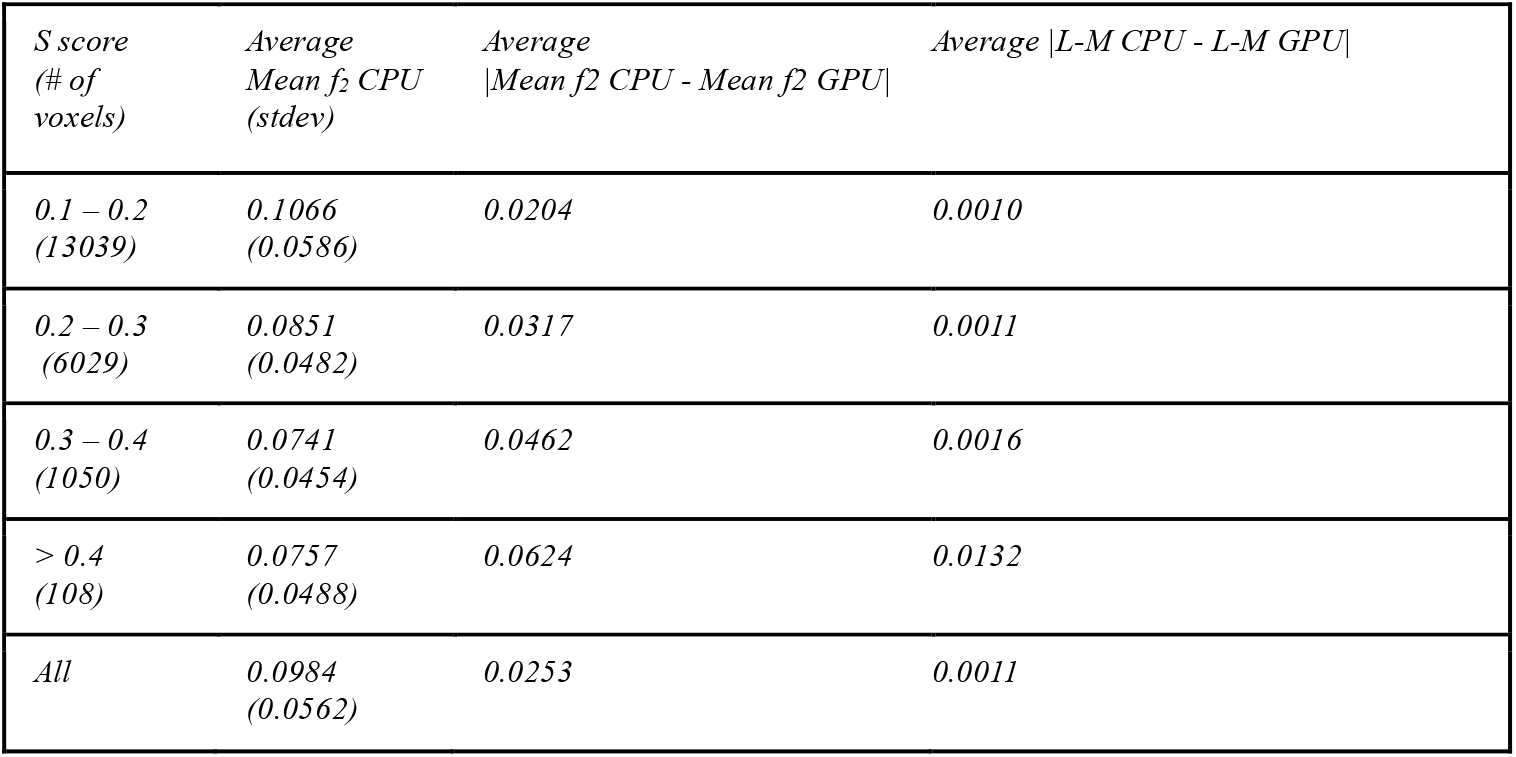
Significantly different *f*_2_ distributions where mean *f*_2_ in CPU or GPU > 0.05: for each S-score range, averaged mean *f*_2_ of CPU distributions, averaged absolute difference in mean *f*_2_ and averaged absolute difference in L-M initialization are tabulated

| *S score*<br>(# of<br>voxels) | *Average*<br>*Mean $f_2$ CPU*<br>(*stdev*) | *Average*<br>*\|Mean $f_2$ CPU - Mean $f_2$ GPU\|* | *Average \|L-M CPU - L-M GPU\|* |
| --- | --- | --- | --- |
| *0.1 – 0.2*<br>(13039) | *0.1066*<br>(*0.0586*) | *0.0204* | *0.0010* |
| *0.2 – 0.3*<br>(6029) | *0.0851*<br>(*0.0482*) | *0.0317* | *0.0011* |
| *0.3 – 0.4*<br>(1050) | *0.0741*<br>(*0.0454*) | *0.0462* | *0.0016* |
| *> 0.4*<br>(108) | *0.0757*<br>(*0.0488*) | *0.0624* | *0.0132* |
| *All* | *0.0984*<br>(*0.0562*) | *0.0253* | *0.0011* |

This was the same threshold chosen by (Behrens et al. 2007) when looking for secondary fibre orientations supported by ARD (Fig. 5). Here, most significantly different voxels were localized in grey/white matter junctions. Some were sparsely found bilaterally within identifiable structures such as corpus callosum, corona radiata, internal capsule, anterior and posterior thalamic radiations.

**Figure 5:**
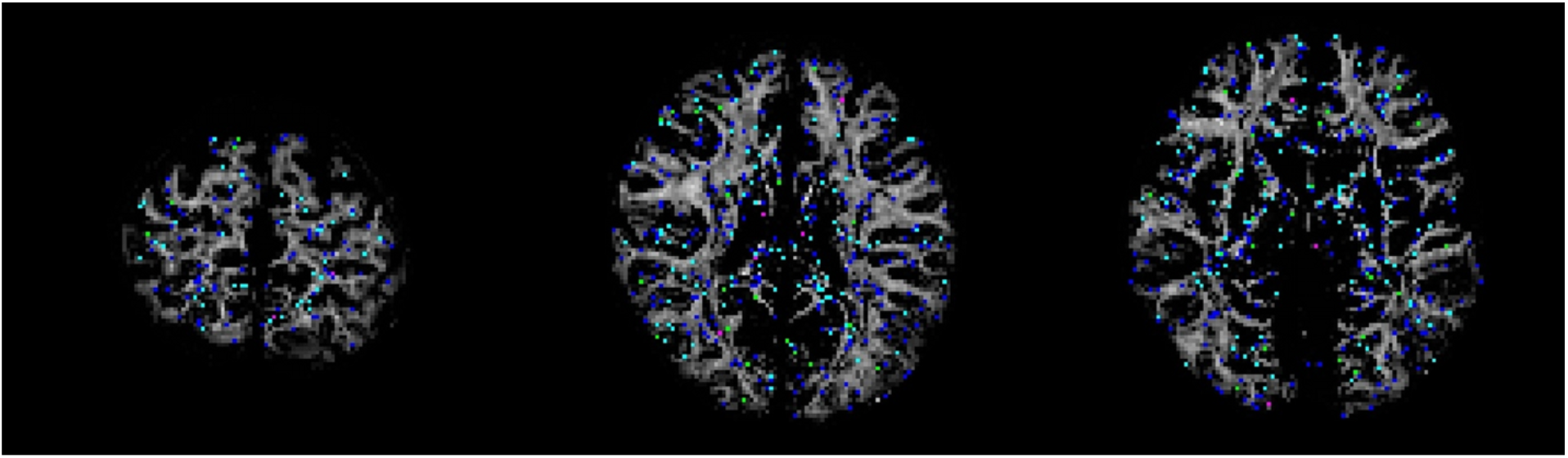
Significantly different *f*_2_ with mean *f*_2_ in CPU or GPU > 0.05. S-score ranges are in: 0.1-0.2=blue, 0.2-0.3=light-blue, 0.3-0.4=green, > 0.4=magenta

The majority of significant *f*_2_ distribution differences had S-scores < 0.3 (19068 out of 20226). One example of a voxel exhibiting a large PDF difference in *f*_2_ is depicted in Fig. 6, where S-score = 0.625 and is the same location where the largest S-score was found for *f*_1_ distribution.

**Figure 6:**
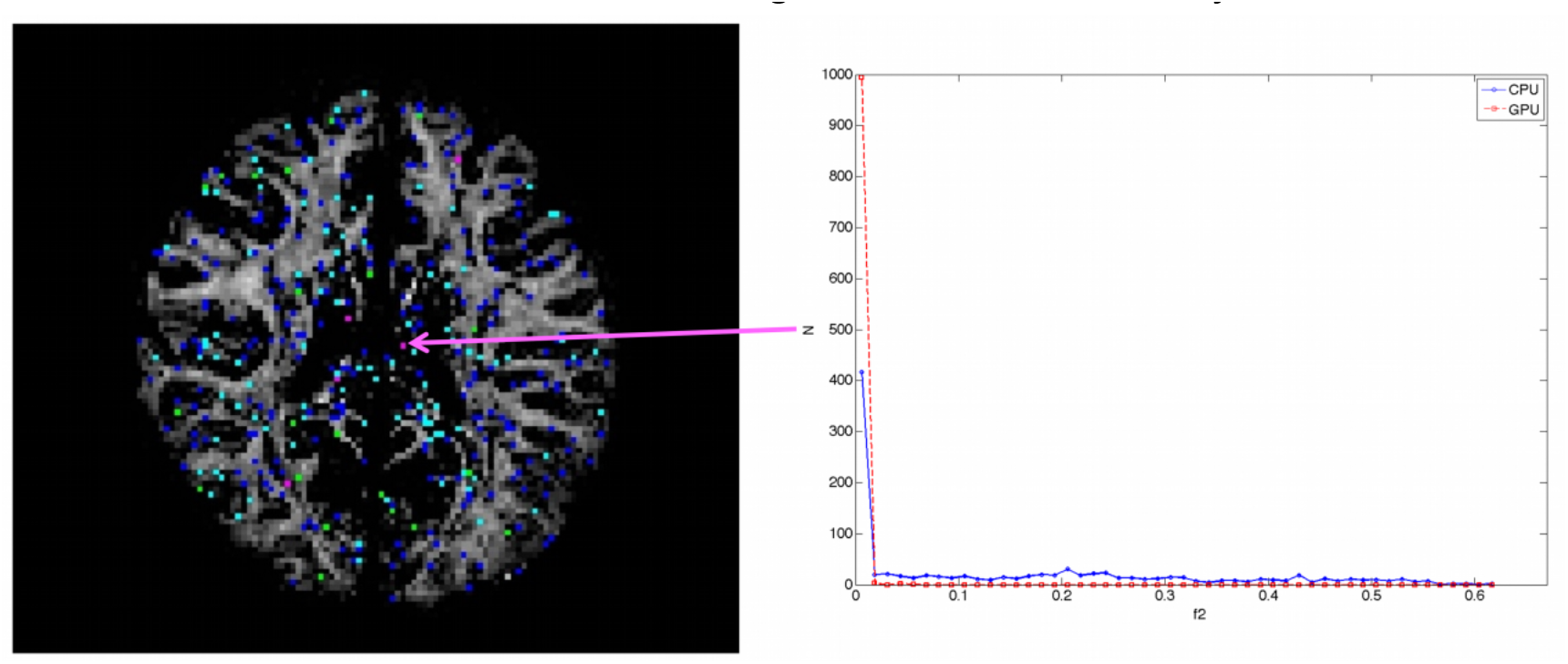
Example distribution shape differences in *f*_2_. Red squares with dotted line denotes GPU samples and blue circles with solid line denotes CPU samples

Here, the distribution shows a sharp peak with low variance near *f*_2_ = 0 in the GPU distribution. In the CPU, there is a smaller peak near *f*_2_ = 0 with a larger variance that spans from 0 to 0.6. The CPU L-M initialization step estimated *f*_2_ = 0.48 averaged across 20 trials while GPU L-M initialization step had *f*_2_ = 0 across 20 trials. Similar to *f*_1_, larger average L-M differences were found for larger S ranges. After adjusting for *f*_2_>*f*_1_ samples, 1303 voxels were no longer significantly different, and of these voxels, 1096 were in areas with mean *f*_2_ from CPU or GPU greater than or equal to 0.05.

### D. ϕ_1_ and θ_1_

196081 out of 436738 total brain voxels had significantly different ϕ_1_ or θ_1_ distributions. Significantly different distributions were localized predominantly in areas of grey matter (43%) and cerebrospinal fluid (37%). They were also found in the white matter (20%), with some key white matter structures such as corpus callosum, internal capsules, corona radiata and anterior and posterior thalamic radiations containing significantly different distributions (Fig. 7).

**Figure 7:**
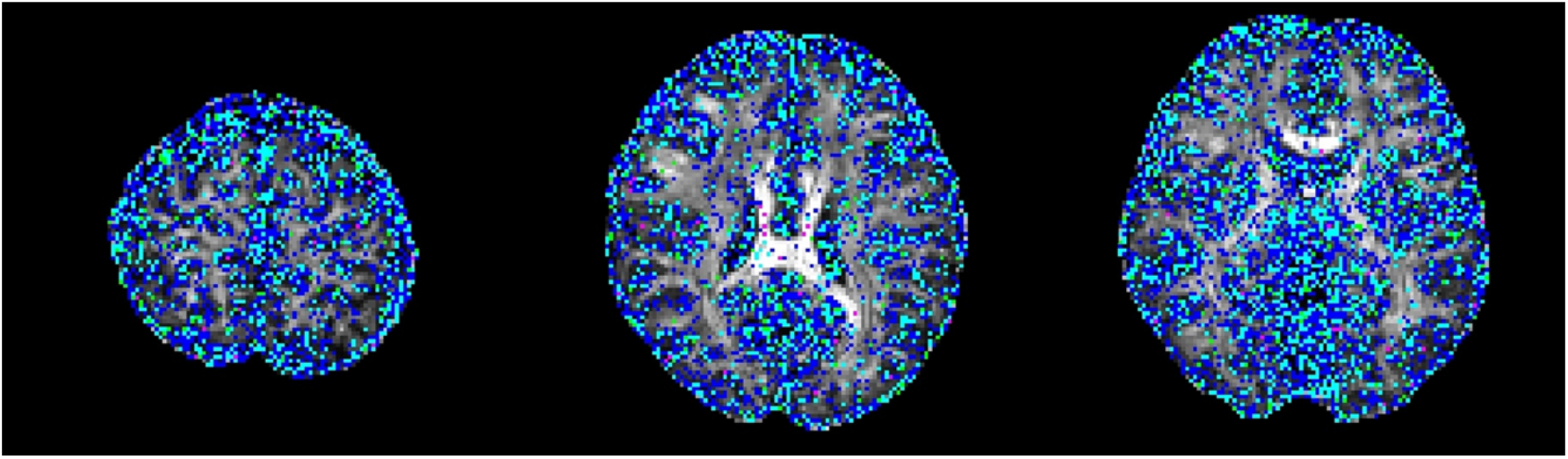
Significantly different ϕ_1_ and θ_1_ distributions with S-ranges in 0.1-0.2 (Blue), 0.2-0.3 (light-blue), 0.3-0.4 (green), > 0.4 (magenta). Maximum S-score between ϕ_1_ and θ_1_ was used to categorize each location into different range

Mean and median angle differences, and average 95^th^ percentile CAUs for each S-score range are tabulated in Table 3.

**Table 3:** Significantly different ϕ_1_, θ_1_: for each S-score range, average of mean PDD difference, standard deviation of mean PDD difference, median of mean PDD difference and 95^th^-percentile cone of angular uncertainty are tabulated.

| *S score*<br>*(# of*<br>*voxels)* | *Average*<br>*Δmean*<br>*PDD* | *Stdev*<br>*Δmean*<br>*PDD* | *Median*<br>*Δmean*<br>*PDD* | *Mean 95<sup>th</sup>-percentile*<br>*CAU* |  |
| --- | --- | --- | --- | --- | --- |
|  |  |  |  | *CPU* | *GPU* |
| *0.1 – 0.2*<br>*(113904)* | *2.146°* | *4.061°* | *1.047°* | *50.841°* | *50.818°* |
| *0.2 – 0.3*<br>*(68528)* | *2.193°* | *4.122°* | *1.080°* | *55.860°* | *55.845°* |
| *0.3 – 0.4*<br>*(12098)* | *2.306°* | *4.584°* | *1.086°* | *58.358°* | *58.448°* |
| *> 0.4*<br>*(1551)* | *2.529°* | *6.394°* | *1.008°* | *57.702°* | *57.346°* |
| *All* | *2.175°* | *4.140°* | *1.061°* | *53.113°* | *53.097°* |

Again, the majority of these voxels have S < 0.3 (182432 out of 196081). Mean difference in angles of principle diffusion directions in all significantly different voxels was 2.175° (stdev = 4.140°) while the median difference was 1.061°. In all significantly different ϕ_1_ and θ_1_ voxels, the average angular difference between the 95^th^ percentile CAUs for CPU and GPU is 0.016° (CPU 53.113°; GPU 53.097°; see Table 3). Because ϕ_1_ and θ_1_ parameters are more meaningful in white-matter where anisotropy is higher, angular differences and 95^th^ percentile CAUs for each S-score range in white-matter only are tabulated in Table 4.

**Table 4:** Significantly different ϕ_1_, θ_1_ with in the white-matter: for each S-score range, average of mean PDD difference, standard deviation of mean PDD difference, median of mean PDD difference and 95^th^-percentile cone of angular uncertainty are tabulated

| *S score*<br>*(# of*<br>*voxels)* | *Average*<br>*Δmean*<br>*PDD* | *Stdev*<br>*Δmean*<br>*PDD* | *Median*<br>*Δmean*<br>*PDD* | *Average 95<sup>th</sup>-percentile*<br>*CAU* |  |
| --- | --- | --- | --- | --- | --- |
|  |  |  |  | *CPU* | *GPU* |
| *0.1 – 0.2*<br>*(20077)* | *2.276°* | *3.150°* | *1.300°* | *38.839°* | *38.754°* |
| *0.2 – 0.3*<br>*(9274)* | *3.382°* | *4.884°* | *1.606°* | *45.891°* | *45.950°* |
| *0.3 – 0.4*<br>*(1372)* | *4.660°* | *7.148°* | *1.672°* | *49.944°* | *49.905°* |
| *> 0.4*<br>*(218)* | *7.140°* | *13.658°* | *1.160°* | *40.633°* | *40.171°* |
| *All* | *2.748°* | *4.210°* | *1.394°* | *41.458°* | *41.415°* |

Overall, higher average difference in mean PDD and lower CAUs were found in significantly different voxels confined to the white-matter. The effect of distribution shape difference on diffusion direction is illustrated in Fig 8. The depicted distributions came from a voxel located posteriorly in the genu of the right internal capsule with S score = 0.55. ϕ_1_ distributions had two notable peaks but there was a difference in height between CPU and GPU. θ_1_ distributions had different number and locations of peak values between CPU and GPU, with the GPU distribution having evenly split peaks in two locations. The resulting mean directions from these distributions differed as depicted in the 3D-plot of Fig 8.

**Figure 8:**
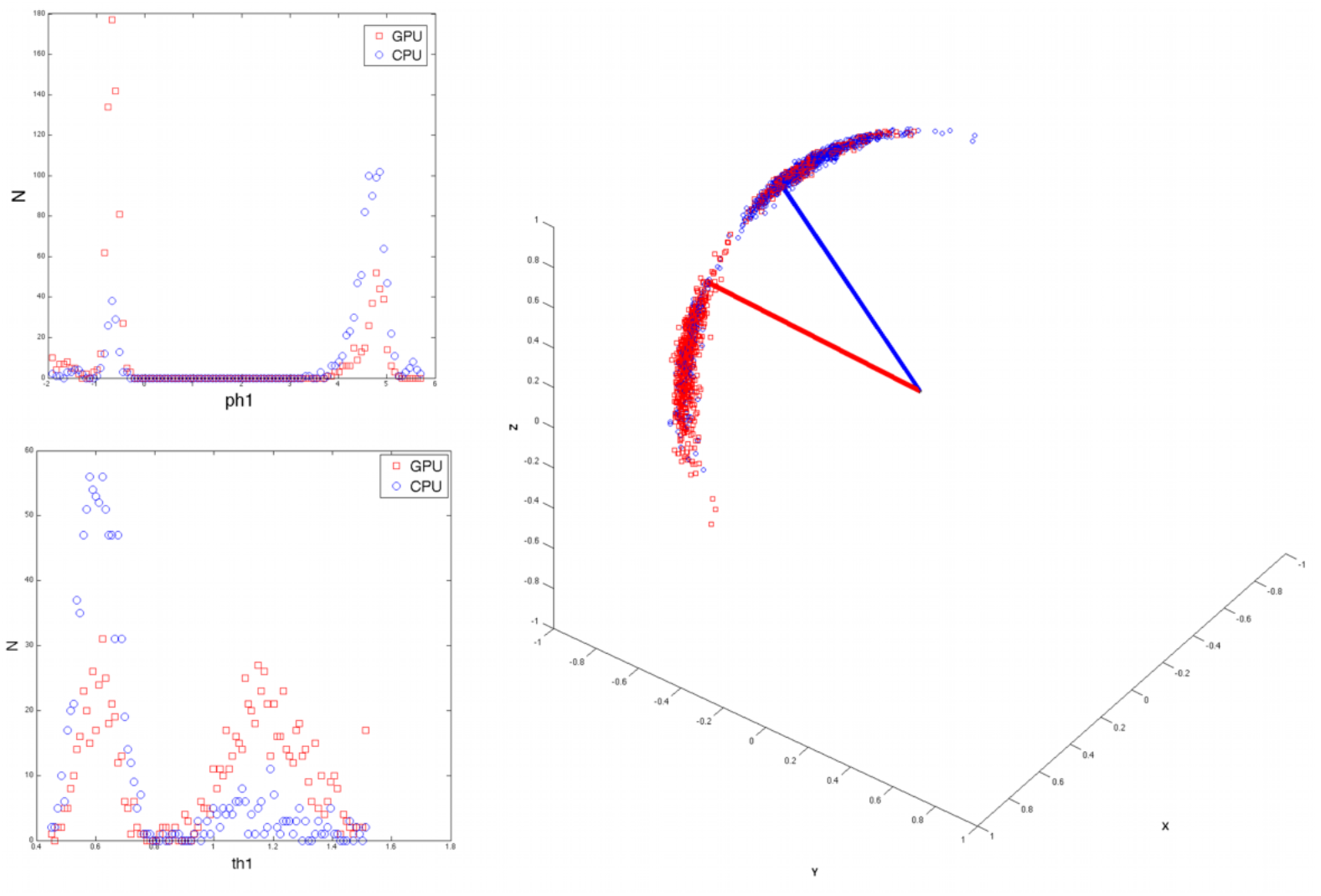
Distributions of ϕ_1_ and θ_1_ that are significantly different were derived from one representative voxel with a particularly high S-score of 0.55. The voxel was located within the genu of the right internal capsule, with voxel coordinates 79, 80, 46. Red squares denote GPU samples while blue circles denote CPU samples. Blue solid line: CPU ϕ_1_, θ_1_ mean direction, Red solid line: GPU ϕ_1_, θ_1_ mean direction. Difference in peak locations and heights result in difference in mean diffusion direction for CPU and GPU

There were no significant differences in *f*_1_ and *f*_2_ in this location, and mean CPU *f*_1_ and *f*_2_ values were 0.411 and 0.351 respectively, signifying that it was suitable for modeling two different fibre orientations with higher anisotropy in this location. Adjusting for f_2_ > f_1_, 33133 voxels became not significantly different and of these, 11043 were in the white-matter.

### E. ϕ_2_ and θ_2_

223309 out of 436738 total brain voxels had significantly different ϕ_2_ or θ_2_ distributions. Significantly different distributions were localized in grey matter (45%), cerebrospinal fluid (25%) and white matter structures (30%) such as corpus callosum, corona radiata, internal capsule, and the anterior and posterior thalamic radiations (Fig 9). Mean and median angle differences along with 95^th^ percentile CAUs for each S-score range are tabulated in Table 5. Again, most voxels have S-scores < 0.3 (212651 out of 223309).

**Figure 9:**
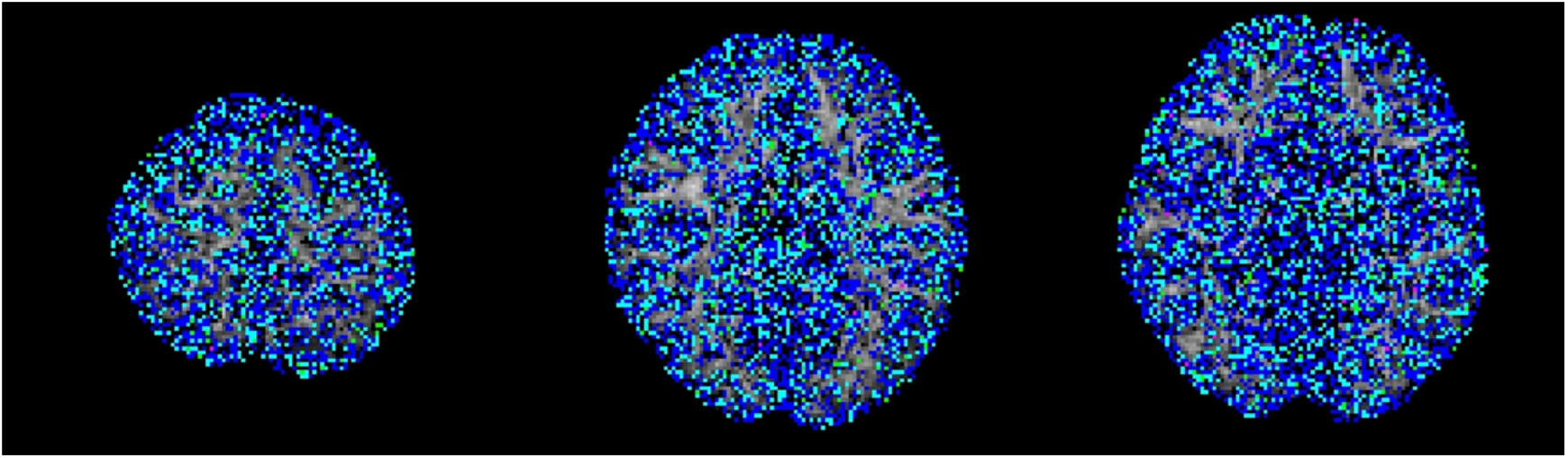
Significantly different ⍰2 and ⍰2 distributions with S-ranges in 0.1-0.2 (Blue), 0.2-0.3 (light-blue), 0.3-0.4 (green), > 0.4 (magenta). Maximum S-score between ⍰2 and ⍰2 was used to categorize each location into different range

**Table 5:** Significantly different ϕ_2_, θ_2_: for each S-score range, average of mean PDD difference, standard deviation of mean PDD difference, median of mean PDD difference and 95^th^-percentile cone of angular uncertainty are tabulated.

| $S$ score<br>(# of<br>voxels) | Average<br>Δmean<br>PDD | Stdev<br>Δmean<br>PDD | Median<br>Δmean<br>PDD | Average 95<sup>th</sup>-percentile<br>CAU | |
| --- | --- | --- | --- | --- | --- |
|  |  |  |  | CPU | GPU |
| 0.1 – 0.2<br>(139823) | 37.019° | 29.345° | 34.518° | 83.236° | 83.229° |
| 0.2 – 0.3<br>(72828) | 35.912° | 29.197° | 31.690° | 84.095° | 84.095° |
| 0.3 – 0.4<br>(9910) | 37.537° | 29.008° | 35.257° | 84.098° | 84.151° |
| > 0.4<br>(748) | 41.154° | 29.247° | 43.285° | 83.276° | 83.359° |
| All | 36.695° | 29.288° | 33.656° | 83.554° | 83.553° |

Overall mean difference in principle directions is 36.695° with median difference of 29.288°. Average angular uncertainty is 83.554° and 83.553° for CPU and GPU, respectively. With ϕ_2_ and θ_2_, it is more meaningful to focus on white matter and areas where *f*_2_ > 0.05 (i.e. where ARD has deemed appropriate to fit a second fibre orientation). Mean PDD difference and 95^th^-percentile CAUs in the white-matter and f_2_ > 0.05 for each S-score range are tabulated in Table 6. Like ϕ_1_ and θ_1_, the CAUs were lower when focusing in on the white matter region. Also, the difference in mean PDD was lower for each S-score range. Distribution shapes of ϕ_2_, θ_2_ samples and the resulting distribution of principle directions in the same location as ϕ_1_, θ_1_ in the genu of the right internal capsule are depicted in Fig 10. There was similarity in direction distribution between [ϕ_1_, θ_1_] and [ϕ_2_, θ_2_] while the latter had more anti-parallel directions included in the distribution. Also, mean directions of CPU and GPU were roughly reversed in [ϕ_2_, θ_2_] compared to [ϕ_1_, θ_1_], showing the difference in how CPU and GPU labelled the underlying multiple fibre orientations. Mean CPU [ϕ_1_, θ_1_] direction and mean GPU [ϕ_2_, θ_2_] direction had a difference of 2.7465°, while mean CPU [ϕ_2_, θ_2_] direction and mean GPU [ϕ_1_, θ_1_] direction had a difference of 18.4179°. After swapping the orientation samples with *f*_2_ >*f_1_*, underlying fibre orientations in this particular voxel became better aligned between CPU and GPU (Fig. 11).

**Table 6:**
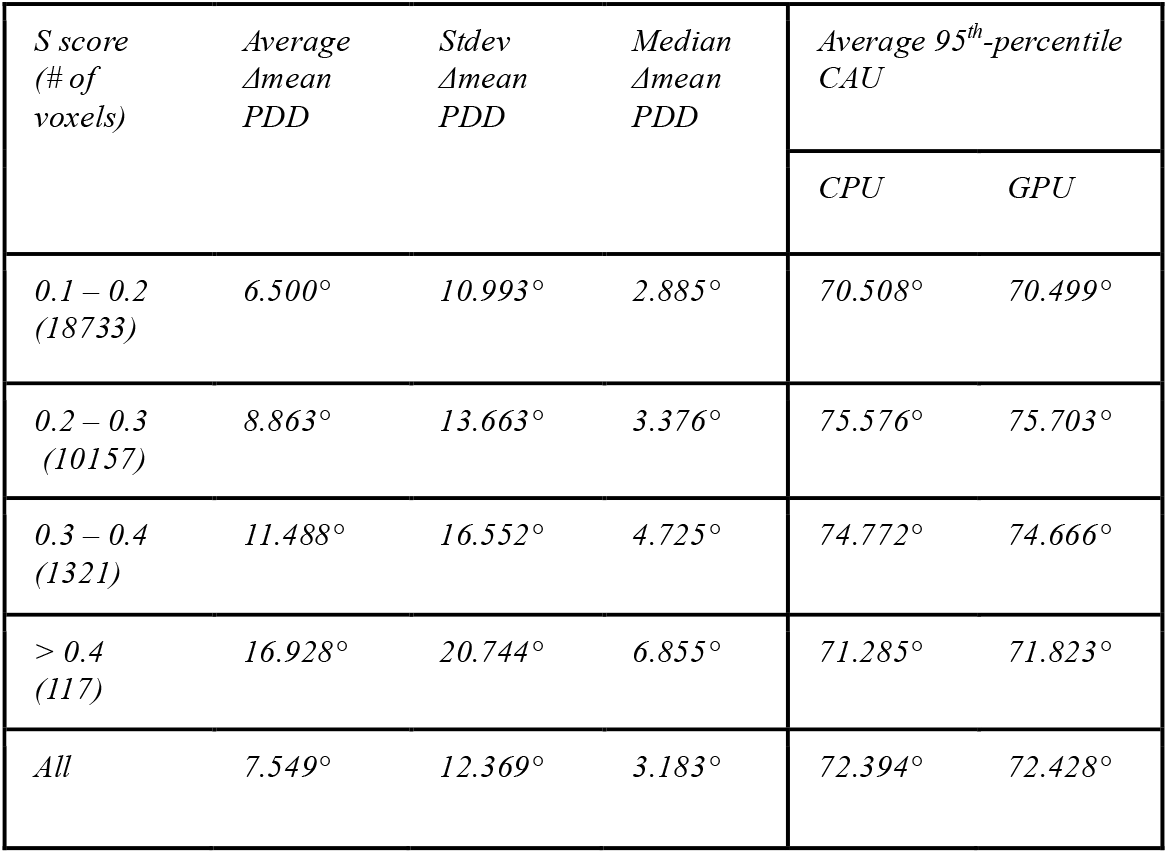
Significantly different ϕ_2_, θ_2_ in the white-matter: for each S-score range, average of mean PDD difference, standard deviation of mean PDD difference, median of mean PDD difference and 95^th^-percentile cone of angular uncertainty are tabulated

| $S$ score<br>(# of<br>voxels) | Average<br>Δmean<br>PDD | Stdev<br>Δmean<br>PDD | Median<br>Δmean<br>PDD | Average 95<sup>th</sup>-percentile<br>CAU | |
| --- | --- | --- | --- | --- | --- |
|  |  |  |  | CPU | GPU |
| 0.1 – 0.2<br>(18733) | 6.500° | 10.993° | 2.885° | 70.508° | 70.499° |
| 0.2 – 0.3<br>(10157) | 8.863° | 13.663° | 3.376° | 75.576° | 75.703° |
| 0.3 – 0.4<br>(1321) | 11.488° | 16.552° | 4.725° | 74.772° | 74.666° |
| > 0.4<br>(117) | 16.928° | 20.744° | 6.855° | 71.285° | 71.823° |
| All | 7.549° | 12.369° | 3.183° | 72.394° | 72.428° |

**Figure 10:**
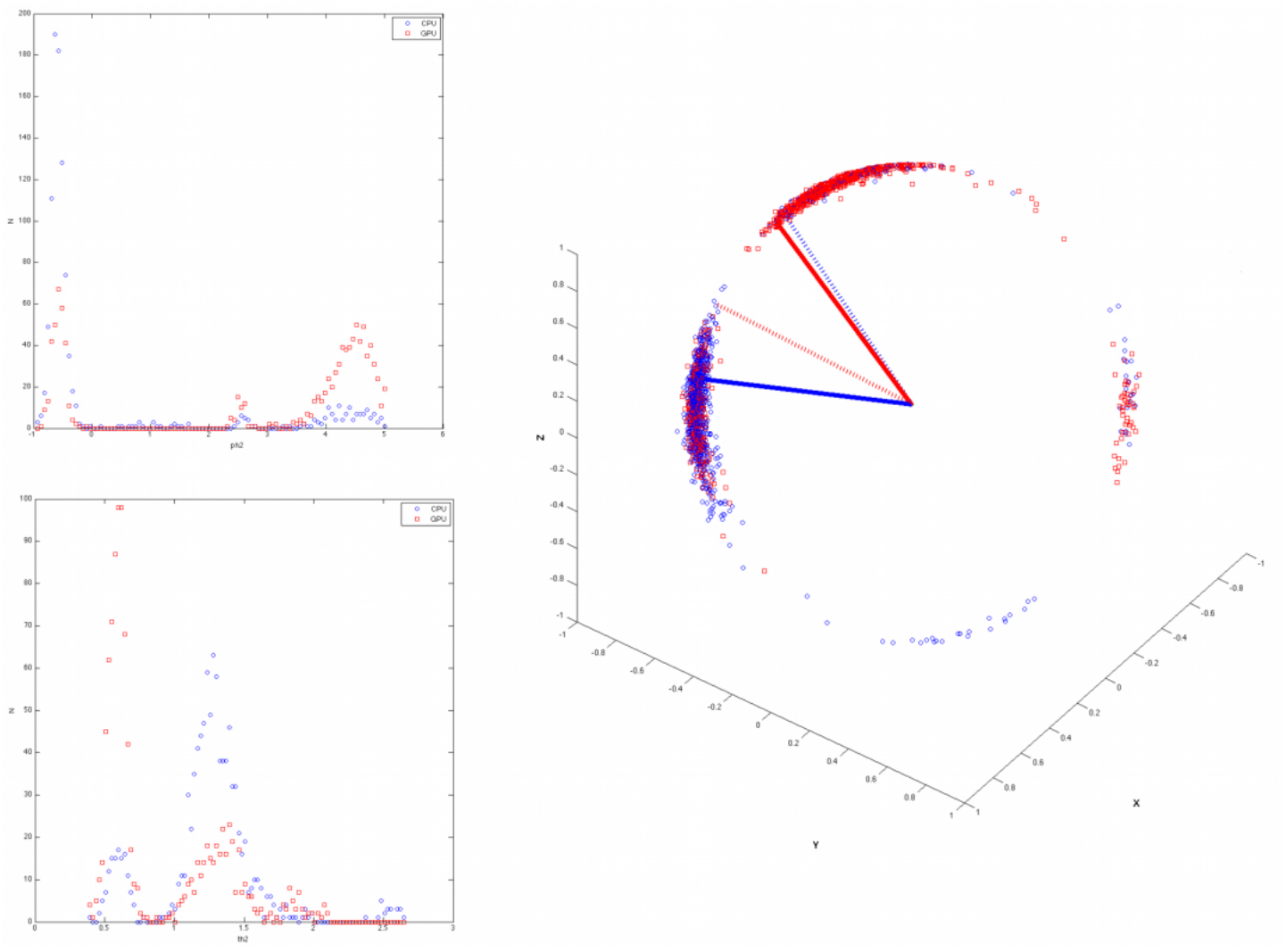
Example ϕ_2_,θ_2_ distributions from the same voxel depicted in Figure 8, within the genu of the right internal capsule. 3D-plot depicts the distribution of directions derived from ϕ_2_,θ_2_ samples (Blue circle: CPU ϕ_2_,θ_2_ samples, Red square: GPU ϕ_2_,θ_2_ samples) and mean directions (Blue solid line: CPU ϕ_2_,θ_2_ mean direction, Red solid line: GPU ϕ_2_,θ_2_ mean direction, Blue dotted line: CPU ϕ_1_,θ_1_ mean direction, Red dotted line: GPU ϕ_1_,θ_1_ mean direction)

**Figure 11:**
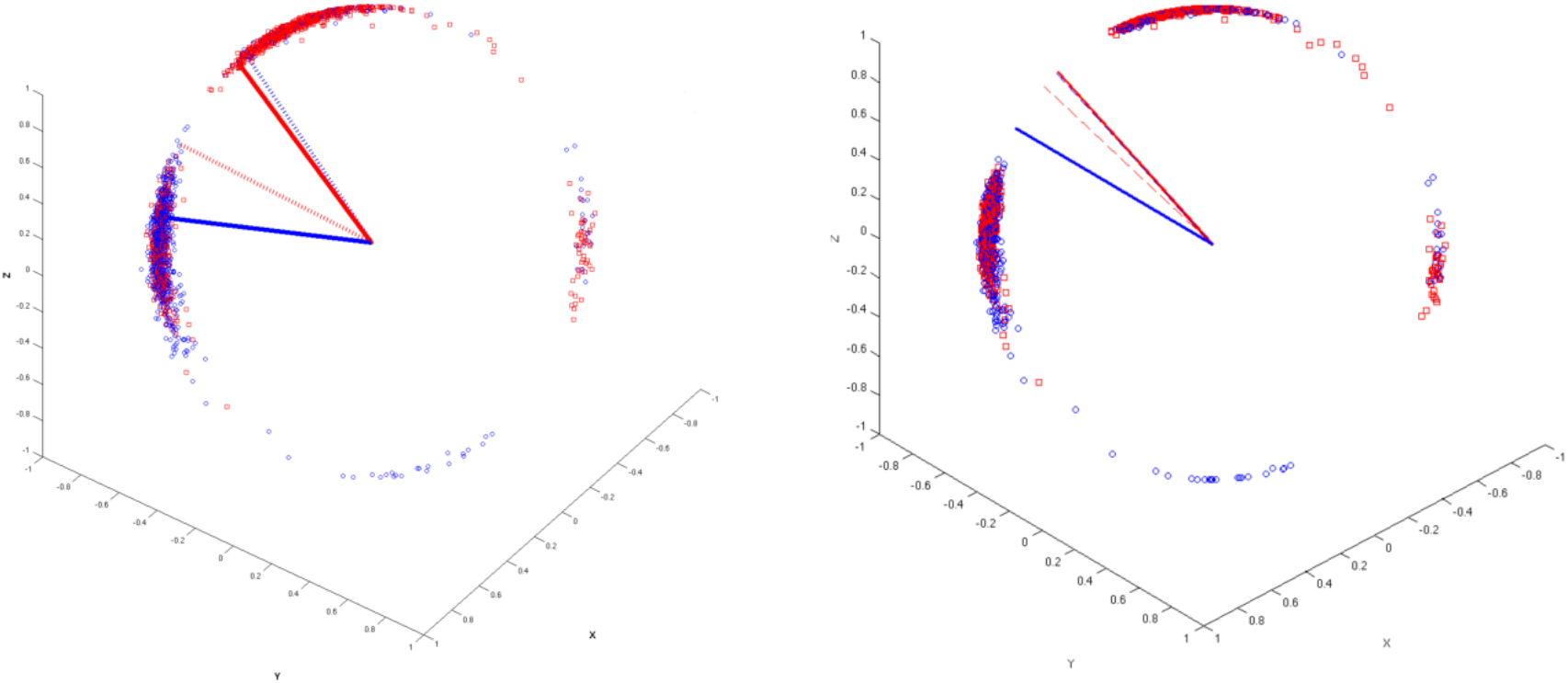
Effect of swapping samples where *f_2_* > *f_1_*: Distribution of directions derived from unswapped ϕ_2_, θ_2_ samples (Left, Blue circle: CPU ϕ_2_, θ_2_ samples, Red square: GPU ϕ_2_, θ_2_ samples) and mean directions (Blue solid line: CPU ϕ_2_, θ_2_ mean direction, Red solid line: GPU ϕ_2_, θ_2_ mean direction, Blue dotted line: CPU ϕ_1_, θ_1_ mean direction, Red dotted line: GPU ϕ_1_, θ_1_ mean direction). Right is swapped samples of directions. Difference in mean principal direction becomes lower when samples are swapped.

## Discussions and Conclusions

A total of 2620428 pairs of distributions were created and compared across the whole brain. 74% of those distributions showed no significant difference between CPU and GPU. Of the significantly different distributions, 44% were localized in grey matter, 31% in cerebrospinal fluid, and 25% in white matter, localized within the corpus callosum in the midline, and bilaterally within the corona radiata, internal capsule, and the anterior and posterior thalamic radiations.

Significantly different *f*_1_ and *f*_2_ distributions in a prominent white-matter structure, such as the body of corpus callosum in the midline as displayed in Fig. 3 and Fig. 6, have been noted with more than half of the samples produced by CPU and GPU differing in value. In general, the corpus callosum contains a well-defined fibre bundle in the Left-to-Right orientation and thus we expected a higher *f*_1_ with lower uncertainty while *f*_2_ expected be estimated closer to 0. To understand where the differences were coming from in this particular voxel, we explored a synthetic ball-and-stick data generated using Dipy 0.16.0 (Garyfallidis et al. 2014), to emulate a single-voxel that is similar in diffusion characteristics as the corpus callosum, and compare the *bedpostx* and *bedpostx_gpu* output to synthetic ground-truth. This allowed the operation orders between CPU and GPU to be kept identical (i.e. single-voxel serial order equals single-voxel parallel order). Here, mean values of all parameter estimates from CPU and GPU were within 1% of synthetic ground-truth values, but *f_1_, f_2_* and *ϕ_2_* distributions between CPU and GPU were found to be with significantly different shapes. This suggests that when operation orders are kept the same (e.g. single-voxel data), difference in distribution shape is minimized between the CPU and GPU, and both algorithms estimated parameters closer to ground-truth. Since the corpus callosum in human brain would generally be found near the centre of the whole brain diffusion data, the relative operation order at which this voxel gets processed between the CPU and GPU would have differed greatly (earlier in GPU vs. later in CPU), and this may explain why significant difference was observed even in a prominent structure such as the corpus callosum.

We also examined the initialization stages of L-M fit and noted that the results differed between CPU and GPU. For the L-M fit algorithm between CPU and GPU, the difference in operation order of when a particular voxel gets initialized is less impactful since L-M fit simply minimizes sum-of-squared residuals (Hernandez-Fernandez et al. 2013). However, the different CPU and GPU-CUDA math libraries can result in different initialization values. Although GPU-CUDA math operations are double-precision capable (Whitehead and Fit-Florea 2011), increased peak performance (e.g. higher speed-up) is found when single-precision operations are used in their place which may differ in precision compared to CPU math operations (Laguna et al. 2019). Our findings from table 1 and table 2 show that larger L-M initialization difference at the start of MCMC sampling results in larger S-scores in significantly different distributions. This suggests that differences in PDF samples appeared to be stemming from a combination of the following: 1) differing starting points after the L-M fit, 2) differing operation order and 3) difference in math library. It is also observed that the L-M initialization difference maps (Fig. 1) spatially resemble the inverse of a typical SNR map from a multi-channel MR head-coil (Gruber et al. 2018; Wiggins et al, 2006). Though we did not directly compare the SNR map of this dataset with L-M differences, we note and speculate that the larger difference in L-M initialization values between CPU and GPU that are mostly found near the inner most structure of the brain could be possible due to less SNR typically found in this region of the brain due to the head-coil SNR profile. This could also possibly explain why large difference was even noted in prominent white matter such as body of corpus callosum: the scan orientation is such that this structure is farther away from the head-coil elements.

We found that S-scores of significantly different distributions were no greater than 0.3 for 94% of significantly different distributions, i.e. 30% or less samples caused the difference. Distributional shape differences were characterized by: a) peak height differences for fibre fractions and b) number of peaks and peak value differences for diffusion direction angles. Larger difference in shape resulted in larger difference in mean values or principle diffusion direction angles. Mean angular differences in principle diffusion directions were 2.175° and 36.695° for significantly different [ϕ_1_,θ_1_] and [ϕ_2_,θ_2_] respectively. Their 95^th^ percentile CAUs were 53.1° and 83.5° respectively. We see the larger CAUs for [ϕ_2_,θ_2_] coming from angle samples that are antiparallel to each other and mislabeled angle samples that may be representing [ϕ_1_,θ_1_] sub-fibre population, thereby increasing the average uncertainty in that location. Algorithms such as *probtrackx* does tract streamlining with tract propagation constraints that propagate streamlines smoothly, and avoid internal looping or sharp turns. This is achieved by treating antiparallel angles as the same (i.e. multiplying antiparallel angles by −1 prior to propagation), and sampling from fibre-population that has minimal angular difference from previous propagation direction. These constraints would effectively allow consistent tract streamlines to be produced from CPUs and GPUs, despite the difference in PDF distributional shapes in the PDD angles. We observed that CPU can produce mean diffusion directions in [ϕ_1_,θ_1_] that are similar to GPU’s mean diffusion directions in [ϕ_2_,θ_2_] and vice-versa (see Fig. 10). Previously, Jbabdi et al. have reported similar inconsistency in sub-fibre population labeling in bedpostx and they consolidated the orientations of sub-fibres by performing a swaping operation, similar to our method of swapping the angular samples where *f_2_>f_1_* (Jbabdi et al. 2010). In our dataset, swapping of angular samples where *f*_2_>*f*_1_ produced better aligned mean directions between CPU and GPU. We note that one of the limitations of this current work is that there was no investigation into effect of using multi-shell models while doing the comparisons. It is reasonable to suggest that with higher b-values there will be better angular resolutions which might lead to better agreement between CPU and GPU results. The challenge in investigating this would be that the CPU *bedpostx* process will take far too long compared to the GPU as much more data need iterative estimation in a linear fashion, and one would require the use of multi-core High Power Computing resources to do this type of investigation in a reasonable amount of time. Still, b=1000 with a monoexponential model estimation is valuable to investigate between CPU and GPU as most clinical MRI can acquire this type of data with conventional MR system setup and speed-up in *bedpostx* in the GPU can be most effective in this type of front-line clinical setting. Another limitation of this work is that the random-number generator type is not the same between CPU and GPU and thus there is no way of telling how much effect the random number generators have on the differences observed between the two algorithms. The authors have looked at preliminary data where the GPU *bedpostx* algorithm was modified to use the linear-congruential random number generator to obtain the same amount of samples and when compared against the CPU samples, they appeared to have similar amount and magnitude of difference as this current work, which suggests the effect of random number generator in producing differing results would be small. This would then lead us to believe that sample differences are more attributable to difference in implementation of CPU-GPU precision points, math libraries between CPU and GPU-CUDA and more importantly the operation order: L-M initialization then MCMC sequentially v.s. L-M parallel then MCMC parallel. As DWI data are collected with greater amount of gradient directions, combination of different b-values, higher-resolutions, conventional DWI post-processing steps will require more computational resources to finish processing in a reasonable amount of time, and GPUs can offer qualitatively the same results with minimal quantitative difference compared to underlying uncertainty with excellent speed (Hernandez-Fernandez 2019).

In summary, although significant differences were found between outputs of CPU and GPU *bedpostx* parameter distributions, differences may have limited impact upon stochastic tractography with singleshell DTI data: a) differences were observed in only 26% of total distributions; b) differences were sparsely distributed in major tract areas; c) differences in fibre orientations were small compared to background angular uncertainty. The latter appears to arise from antiparallel angles and random assignment of principle directions to sub-fibre populations.

## Abbreviations

CPU: central processing unit
GPU: graphics processing unit
bedpostx: Bayesian estimation of diffusion parameter obtained using sampling techniques
dMRI: diffusion MRI
DWI: diffusion weighted imaging
DTI: diffusion tensor imaging
MCMC: Markov Chain Monte Carlo
ARD: automatic relevance determination
L-M: Levenberg-Marquadt
CAU: cone of angular uncertainty
PDF: probability density function
K-S: Kolmogorov-Smirnov

## Declarations

### Ethics approval, consent to participate and consent for publication

Human MRI data used for this study is from a public repository of The Human Connectome Project. The Human Connectome Project (Principal Investigators: Bruce Rosen, M.D., Ph.D., Martinos Center at Massachusetts General Hospital; Arthur W. Toga, Ph.D., University of California, Los Angeles, Van J. Weeden, MD, Martinos Center at Massachusetts General Hospital) is supported by the National Institute of Dental and Craniofacial Research (NIDCR), the National Institute of Mental Health (NIMH) and the National Institute of Neurological Disorders and Stroke (NINDS). Collectively, the HCP is the result of efforts of co-investigators from the University of California, Los Angeles, Martinos Center for Biomedical Imaging at Massachusetts General Hospital (MGH), Washington University, and the University of Minnesota.

### Competing Interest

The authors declare that they have no conflict of interest.

### Funding

Canadian Foundation for Innovation (Project number: 20494)

## Author’s contribution

DHK: data analysis, compilation of results, drafting of manuscript

LJW: drafting of manuscript

MHF: data analysis, drafting of manuscript

BHB: drafting of manuscript

## Availability of data and material

Diffusion and 3D T1-weighted anatomical scans are available through the MGH-USC Human Connectome Project Image & Data Archive portal (https://ida.loni.usc.edu/). Subject mgh1005 was used for analysis.

Software used for this work is available through FSL (https://fsl.fmrib.ox.ac.uk/fsl/fslwiki/). The specific version of FSL can be obtained through the “fsl-5.0” package in Neurodebian repository (http://neuro.debian.net/pkgs/fsl-complete.html)

## Acknowledgment

DHK, LJW, and BHB would like to acknowledge the support of the Pediatric Neurosciences Program, a joint program of the Divisions of Neurology and Neurosurgery at British Columbia Children’s Hospital and the BC Children’s Hospital Research Institute, University of British Columbia. We would like to thank Mr. Kevin Fitzpatrick, Dr. Jonathan Nakane and Dr. Qing-San Xiang for comments on previous drafts of this manuscript.

## Notes

### Competing Interest Statement

The authors have declared no competing interest.

### Summary of Updates

Abstract, Background and Discussion sections updated with more up-to-date references, wording changes; Author affiliations updated

